# A Microglial Subset at the Tumor-Stroma Interface of Glioma

**DOI:** 10.1101/2020.10.27.357459

**Authors:** Michael D. Caponegro, Ki Oh, Miguel Madeira, Daniel Radin, Nicholas Sterge, Richard A. Moffitt, Stella E. Tsirka

**Affiliations:** Program in Molecular and Cellular Pharmacology, Department of Pharmacological Sciences, Stony Brook University, Stony Brook, NY, USA; Department of Biomedical Informatics, Stony Brook University, Stony Brook, NY, USA; Department of Pathology, Stony Brook University, Stony Brook, NY, USA

**Keywords:** Microglia, GAM, Glioma, Leading edge, CCL2, P2RY12

## Abstract

**Background:** Myeloid involvement in High Grade Gliomas, such as Glioblastoma, has become apparent and detrimental to disease outcomes. There is great interest in characterizing the HGG tumor microenvironment to understand how neoplastic lesions are supported, and to devise novel therapeutic targets. The tumor microenvironment of the central nervous system is unique as it contains neural and specialized glial cells, including the resident myeloid cells, microglia. Glioma-associated microglia and peripherally infiltrating monocytes/macrophages (GAM) accumulate within the neoplastic lesion where they facilitate tumor growth and drive immunosuppression. A longstanding limitation has been the ability to accurately differentiate microglia and macrophage roles in pathology, and identify the consequences of the spatial organization of these cells.

**Results:** Here we characterize the tumor-stroma border and identify peripheral glioma-associated microglia (PGAM) at the tumor leading edge as a unique subpopulation of GAM. Using data mining and analyses of samples derived from both murine and human sources, we show that PGAM exhibit a pro-inflammatory and chemotactic phenotype that is associated with peripheral monocyte recruitment, poorly enhancing radiomic features, and decreased overall survival.

**Conclusions:** PGAM act as a unique subset of GAM, at the tumor-stroma interface, corresponding to disease outcomes. We propose the application of a novel gene signature to identify these cells, and suggest that PGAM constitute a cellular target of the TME.

**Graphical abstract:** 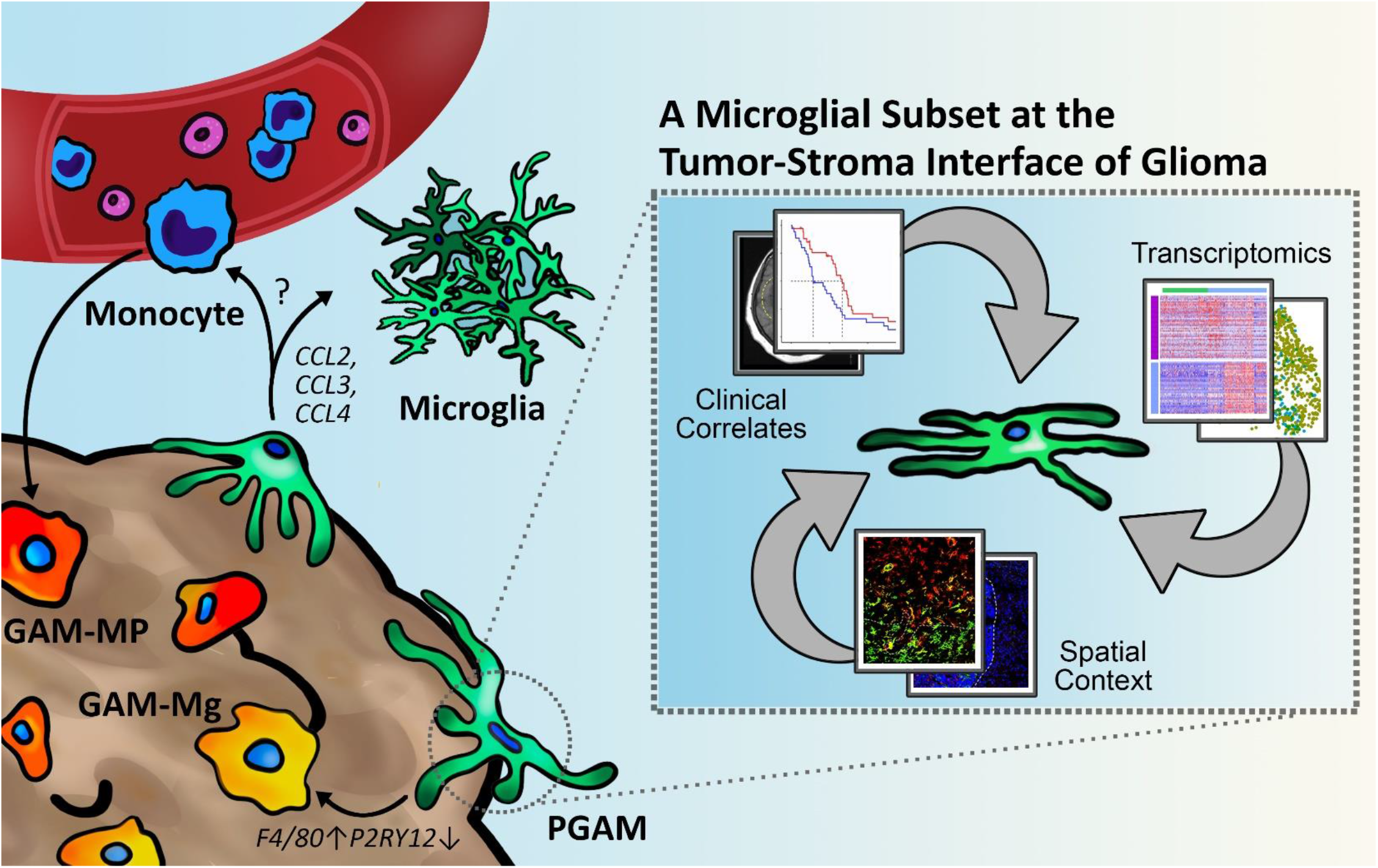

## 1. Introduction

Infiltrating myeloid cells can comprise up to a third of high grade glioma (HGG) lesions^1,2^. As gliomas arise from central nervous system (CNS) tissue, the tumor microenvironment (TME) is unique in that it contains neurons, astrocytes, and the resident myeloid cells, microglia. Microglia act as innate immune hubs of the CNS and influence many neuropathological scenarios. Confounding their roles in disease, inflammation and disruption of the blood-brain barrier result in the recruitment of monocytes/macrophages from the peripheral immune system, which lead to worsened disease outcomes. Despite their distinct cellular ontology^3,4^, microglia and macrophages have been virtually indistinguishable in pathological microenvironments in the CNS, as they express similar biological markers and phenotypic properties. In HGG, it has become well recognized that glioma-associated microglia/macrophages (GAM) are associated with tumor support, therapeutic resistance, and poor survival^5-7^. However, the distinct roles of microglia versus macrophage in HGG pathology is still not fully understood.

Recent application of powerful sequencing technologies, such as single cell and bulk RNA sequencing, have identified key transcripts that differentiate microglia from macrophages, e.g., *P2RY12, TMEM119,* and *OLFML3^8-11^,* and additional comprehensive analyses have been able to parse transcriptional differences between GAM microglia (GAM-Mg) and GAM macrophages (GAM-MP) in the TME^12-15^. Interestingly, these transcriptional identities are paralleled in human and mouse myeloid cells^9,16,17^. As a result, several GAM therapeutic targets have been proposed, although their effectiveness has yet to be determined^18-21^.

An additional complication of GAM transcriptional identities are phenotypic alterations that arise within local TME niches, such as hypoxic cores, necrotic lesions, hyper-proliferative vasculature, and even the expanding tumor margins. Recently, several GAM-Mg genes, including *P2RY12, CX3CR1,* and *SIGLEC8,* were found to associate with the leading edge of glioma^12^. A second theme which has been corroborated across several transcriptional datasets is that GAM-Mg also seem to exhibit a spectrum of activation, i.e. a percentage of cells express relatively high levels of inflammatory markers (*CCL3*, *CCL4, CCL3L1, CCL3l3, IL1A, IL1B, IL8, TNF),* while the remainder resemble homeostatic microglia^10,16,22-24^. While no explanation has emerged from the data, one study suggests that this heightened state of activation may be inherent to age-related microglia phenotypes^23^. Nevertheless, this pro-inflammatory status contradicts the global GAM paradigm, which portrays these populations as universally anti-inflammatory and pro-tumorigenic. Taken together, the lack of phenotypic clarity poses an interesting question regarding GAM-Mg at the leading edge of glioma; are cells of this TME niche unique, and if so, what consequences do their presence have on malignant pathologies?

Until recently, molecular analysis of patient bulk samples have often been confounded by mixed biological samples barring high-resolution analysis of the leading edge and tumor margins. In a recent publication, Darmanis et al sampled matched tumor core and peritumoral tissue from 4 subjects diagnosed with primary Glioblastoma (GB), and performed single-cell RNA sequencing (scRNAseq)^25^. Neoplastic cells of the invading glioma front were found to possess unique transcriptional and genetic profiles, ultimately rendering invasive phenotypes. When myeloid populations were characterized, peritumoral samples were enriched for microglia-specific transcripts, as well as inflammatory transcripts *CCL2, CCL3, CCL4, IL1A, IL1B, TNF, CXCL12,* and *IL6ST*^25^. This finding suggests that the pro-inflammatory and chemotactic signature consistently observed in activated GAM-Mg could be directly resultant from microglia in peripheral CNS tissue, present at the tumor leading edge. However, these peripheral glioma-associated microglia (PGAM) have not been defined or recognized, and their contributions, if any, to HGG pathology are still unknown.

Through comprehensive data mining and integration, and the use of murine *in vivo* models, we demonstrate that PGAM exist as a subset of GAM, and are located at the leading edge niche of HGG. Across the exceedingly large and heterogeneous datasets characterizing microglia in health and disease, we carefully define a transcriptional fingerprint of PGAM. Lastly, we use this information to infer disease outcomes, and suggest potential therapeutic targets of this myeloid compartment in HGG, which heavily contributes to this devastating CNS malignancy.

## 2. Results

### Identification of Peripheral Glioma-Associated Microglia

To begin examining the hypothesis that PGAM are a distinct subset of GAM, we analyzed the myeloid cells of the Darmanis et al. dataset^25^ (Figure 1A). In agreement with published results, our analysis identified a discrete cluster of peripheral myeloid cells (Figure 1B). To further define this population, cells in the myeloid cluster from the ‘peripheral’ tissue sample were selected for by the expression of previously reported microglia-specific genes^9,25^*(TMEM119, P2RY12, GPR34, OLFML3, SLC2A5, SALL1,* and *ADORA3,* Figure 1B). 92.4% of myeloid cells from the peripheral tissue sample were positive for microglia selection criteria. The remaining cells of the myeloid clusters expressed markers for microglia and macrophages, but also included other myeloid-derived markers such as *CD11c* and *CD33,* which in total represent classically-described GAM populations of the tumor core. Expression of microglia-specific transcripts were mapped across the entire myeloid cluster (Figure 1C); notably, *P2RY12* localized almost exclusively to the PGAM group. Supplemental Figure 1 lists the proportion of cells expressing all 7 microglial-specific transcripts, across their respective tumor compartments.

**Figure 1.**
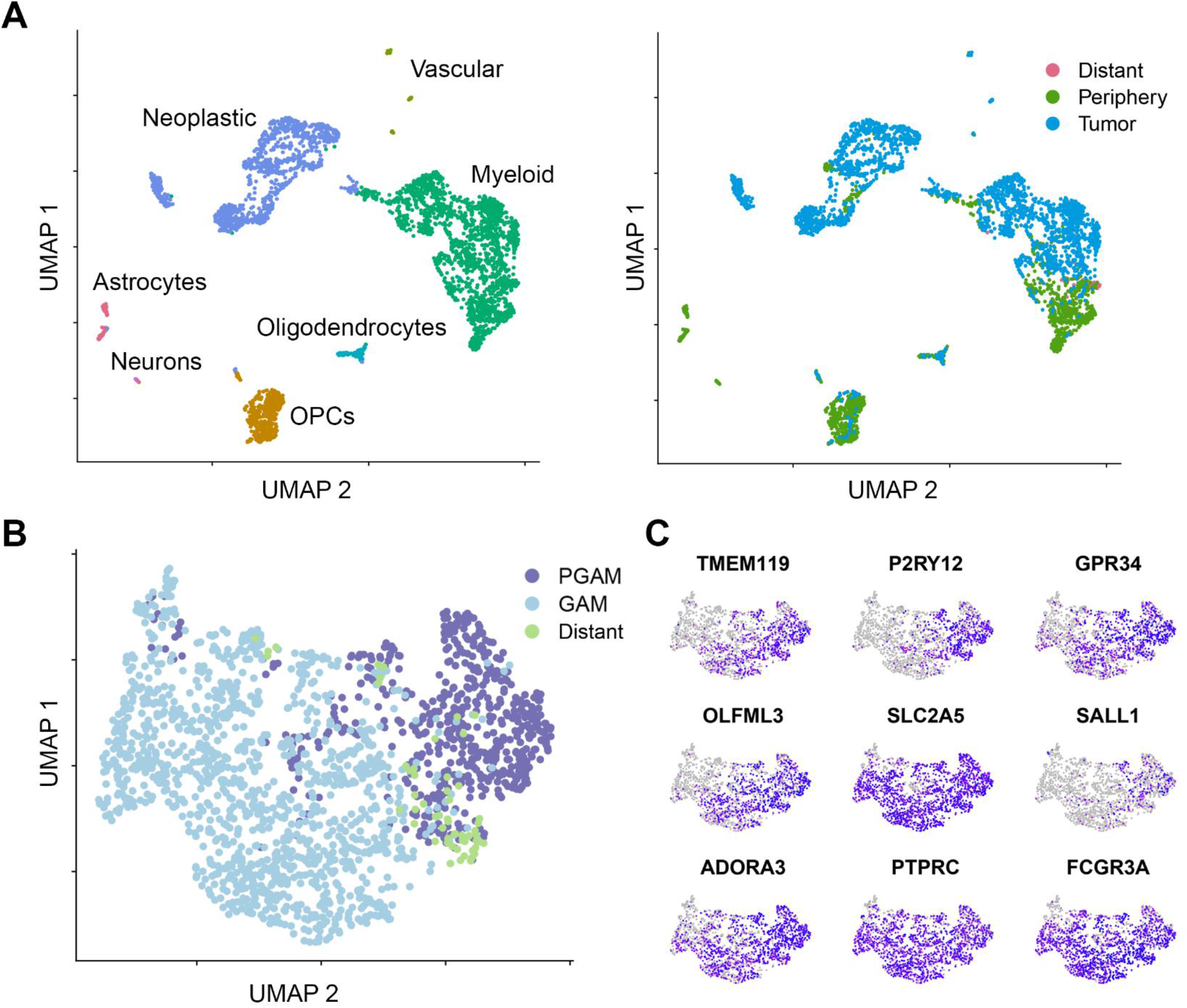
Identification of peripheral glioma-associated microglia. scRNAseq analysis from Darmanis et. al. dataset. **A.** Following dimensional reduction, cells were annotated based on canonical marker expression (left). Myeloid cells segregate with respect to samples from tumor core, or adjacent CNS tissue (right). **B.** Myeloid cells from the peripheral tissue sample were selected for microglial core gene expression (TMEM119, P2RY12, GPR34, OLFML3, SLC2A5, SALL1, and ADORA3), and annotated as PGAM (n = 570). All other myeloid cells were labeled as bona fide GAM (n = 1254). Samples were allowed to re-cluster via UMAP. **C.** Microglial core gene expression plotted across the new myeloid clustering from panel B.

Given that PGAM segregated from bulk tumor GAM in this initial analysis, we sought to determine whether there were transcriptional differences between GAM of the tumor core, and PGAM. Differential expression (DE) analysis identified several biologically relevant transcripts in PGAM. The DE transcripts were ranked and Gene Set Enrichment Analysis (GSEA) was performed^26^. The results from these analyses demonstrated that PGAM were enriched for chemotactic and pro-inflammatory programs, carrying high relative expression of *CCL2, CCL3, CCL4, CCL3L1, CXCL12, CSF1, TNF,* and *IL1B* (Figure 2A). Using the entire scRNAseq dataset, potential cytokine and chemokine receptor-ligand pairs were predicted, which revealed elevated patterns of ligand expression in PGAM, and extensive crosstalk between GAM and PGAM, as well as between the myeloid and neoplastic cells (Figure 2B). These results suggest the existence of a unique subpopulation of microglial cells in GB, which reside in the peritumoral space and exhibit a chemotactic and pro-inflammatory phenotype.

**Figure 2.**
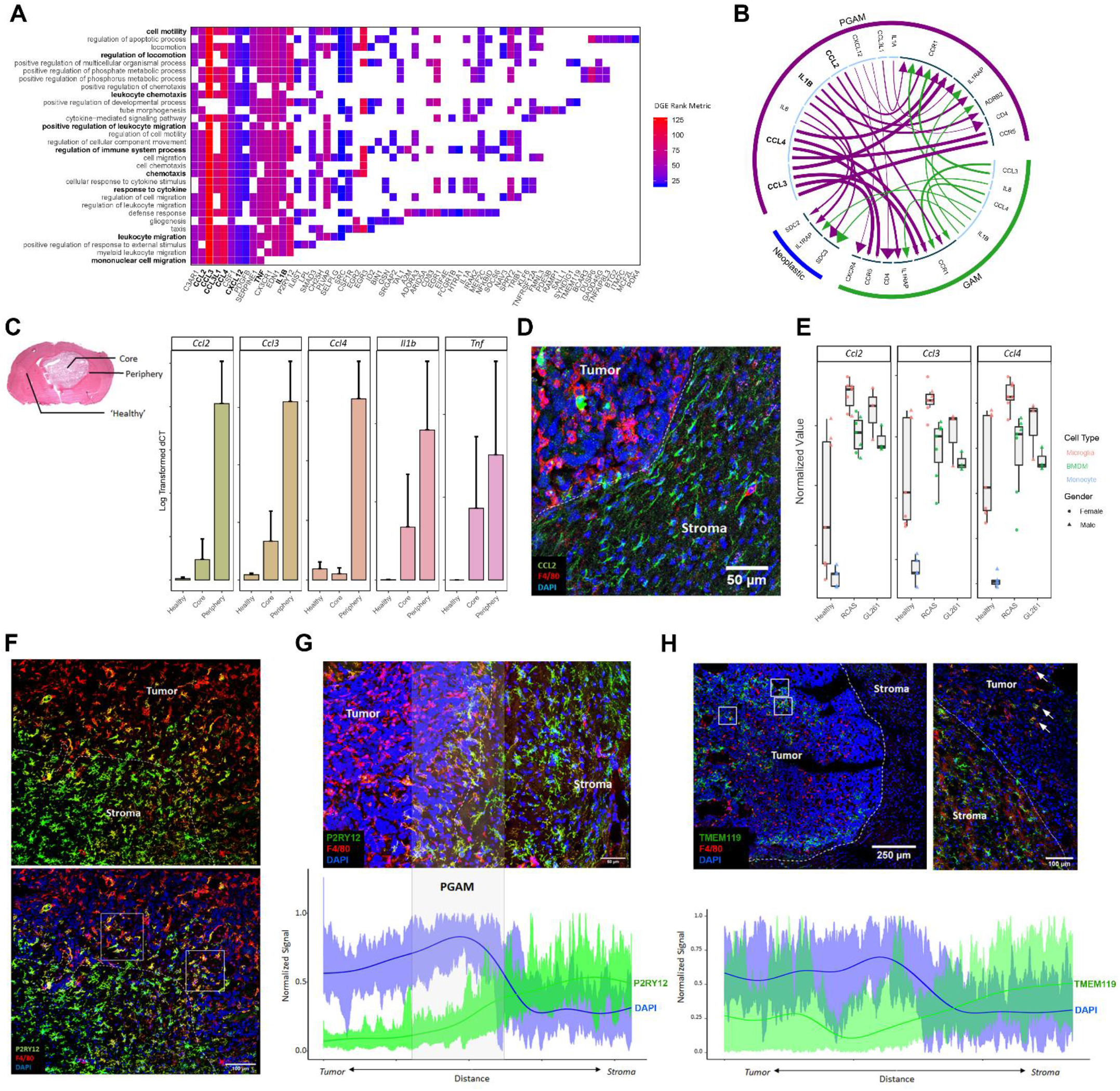
PGAM are chemotactic and pro-inflammatory, and PGAM markers localize at the tumor/stroma interface in glioma. **A.** Transcripts identified from differential gene expression are plotted against their associated GO terms from GSEA pval < 0.01 FDR < 0.05. Axes are sorted such that highly overlapping transcripts and GO terms group together. Tiles are colored by fold change rank metric, relative to tumor core GAM expression. **B.** Using the iTALK package, receptorligand pairs are predicted based on relative expression, between PGAM, GAM, and neoplastic cells. Arrows are colored by cellular population. Arrow lines are scaled by ligand expression, arrow heads are scaled by receptor expression. Inner circle: light blue denotes ligands; dark blue denotes receptor. **C.** Left, representative H&E image of GL261 murine glioma, indicating where contralateral healthy, tumor core, and tumor peripheral tissues were sampled. Right, qRT-PCR log transformed ΔCT values of PGAM markers detected in A, across sample region. n = 2-3. **D.** CCL2^+^ cells populate peritumoral space in GL261 glioma samples. F4/80 is a pan macrophage/microglia marker. n = 3. **E.** bulk RNAseq samples from Bowman et al, and Haage et al were combined, and PGAM markers were plotted against FACS sorted cell types, and murine glioma models GL261 and RCAS. PGAM markers associate with microglia populations, over BMDM and monocytes in both models. n = 3-6.. **F.** P2RY12 staining at the tumor/stroma border in GL261 murine glioma, day 21. White dotted line represents tumor leading edge, white boxes highlight P2RY12^+^F4/80^High^ microglia. Scale bar = 100 μm. **G.** Semiquantitative analysis of spatial distribution of P2RY12 expression across the tumor/stroma interface in glioma, as in F. Spatial signal from imaging technical replicates were scaled and combined, generating an average expression curve. The highly nucleated tumor, marked by high DAPI expression, correlates with the tumor border. n=4. Scale bar = 50 μm. **H.** Semiquantitative analysis of spatial distribution of TMEM119 expression inside the tumor core (left) and across the tumor/stroma interface (right) in glioma, as in G, F. White boxes highlight patches of intratumoral TMEM119^+^F4/80^High^ microglia. White arrows mark TMEM119^+^F4/80^High^ microglia at tumor/stroma interface. n=3. Scale bar = 250 μm and 100 μm.

To validate the chemotactic and pro-inflammatory phenotype observed in peritumoral microglia uncovered from the scRNAseq analysis of the human datasets, we established orthotopic gliomas in immunocompetent C57Bl6 mice, using 2 relevant glioma cell lines (GL261 and KR158B)^27-29^. Gliomas were micro-dissected, and samples of tumor core, periphery, and contralateral hemisphere (representing healthy tissue) were obtained (Figure 2C, left). Quantitative real-time PCR was performed on cDNA libraries generated from the matched tissue samples, and gene expression of top PGAM transcriptional markers were analyzed (Figure 2C). Relative expression of pro-inflammatory markers *Ccl2, Ccl3, Ccl4, Il1b*, and Tnf were all found to be elevated in the peritumoral sample, corroborating the results from DE and GSEA. In a series of recent reports, strong evidence has emerged delineating a CCL2/CCR2-signaling axis in HGG and other solid cancers, where CCL2 expression is correlated with monocyte/macrophage recruitment to the TME, poor patient prognosis, and increased blood flow to the neoplasm^30-33^. Targeting this pathway in preclinical mouse models of HGG has resulted in improved outcomes^1,4,5^. In addition, a recent report has identified that myeloid cells in HGG taken from tumor core and peripheral samples form two signature populations, separated based on levels of *CCL3* and *CCL4* expression, although detailed characterization of populations was not conducted^24^.

Immunofluorescent staining of fixed tissue sections from the murine samples confirmed a high density of CCL2^+^ cells at the tumor/stroma border, compared to cells of the tumor core, in both cell lines used (Figure 2D, Supplemental Figure 2A). To support these results, we utilized recently published FAC-sorted GAM bulk RNAseq datasets, derived from the GL261 and RCAS murine glioma models^8,14^ and found that the expression of *Ccl2*, *Ccl3,* an *Ccl4* were markedly elevated in GAM-Mg over GAM-MP and peripheral blood monocyte populations (Figure 2E). It should be noted however that homeostatic microglia express these genes as well. Indeed, we did find CCL2^+^ cells in contralateral, healthy tissue sections of our samples (Supplemental Figure 2B). Across all cell types in the Darmanis et. al. single cell data set, global levels of *CCL2, CCL3,* and *CCL4* were elevated in the myeloid cell population of the peripheral tissue sample (Supplemental Figure 2C). As myeloid cells comprise the vast majority of non-cancerous cells types found in each patient sample (Supplemental File 1), both the relative levels and absolute magnitude of expression of these molecules are elevated in myeloid cells of the periphery. These data suggest that CCL2 is predominantly expressed by GAM-Mg, spatially maps to the glioma leading edge, and is enriched at the single cell level in the proposed PGAM population. The use of several orthogonal methods corroborates the existence of this unique chemotactic and pro-inflammatory PGAM subpopulation at the tumor/stroma interface in glioma.

Among cell signaling molecules differentially expressed between PGAM and core GAM, *P2RY12* was ranked as the 6^th^ most DE gene, consistent with its role as a microglia-specific marker^9,34^. To investigate the spatial distribution of P2RY12^+^ microglia in glioma, and interrogate its relevance as a potential marker of PGAM, immunostaining for P2RY12 was conducted at different timepoints following glioma inoculation (Figure 2F, Supplemental Figure 3A, B). A distinct double positive population of P2RY12^+^F4/80^High^ cells was visualized exclusively at the tumor/stroma interface, in contrast to P2RY12^+^F4/80^Low^ microglia in the adjacent CNS tissue, and P2RY12^-^F4/80^High^ GAM of the tumor core (Figure 2F, Supplemental Figure 3B, C, D). Semi-quantitative spatial analysis of P2RY12 signal across the tumor/stroma border identified a zone, approximately 200-250μm in width, surrounding the glioma leading edge, harboring double positive P2RY12^+^F4/80^High^ cells (Figure 2G). This pattern is reminiscent of previously observed activation spectrum in GAM-Mg at the single cell transcriptional level.

In contrast, probing for TMEM119 in the same samples as an additional microglia-specific transcript^9^, produced a different staining pattern. TMEM119^+^ cells were identified both within the tumor core as well as along the tumor border (Figure 2H). These results are in agreement with previous reports which describe differential and dynamic expression of microglia-core transcripts during pathological scenarios^35,36^. They also imply that GAM-Mg retain TMEM119 expression after incorporation into the neoplastic lesion, while P2RY12 marks GAM-Mg exclusively across the tumor/stroma interface.

It should be noted that the staining marking these double positive peritumoral microglia was not homogeneously detected around the entire bulk glioma leading edge, but rather was evident in several patches along the invading margin. This finding is consistent with recent reports which identify community-level behaviors along the leading edge of glioma^37,38^. Similarly, cell bodies of mature neurons among neoplastic cells were detected at different areas along the tumor/stroma interface (Supplemental Figure 3E, F).

Together, these results validate the presence of chemotactic/pro-inflammatory PGAM at the tumor invasive margin, offer a microglia-specific cell surface protein as a marker to identify PGAM, and characterize changes in microgliaspecific gene expression with spatial context in HGG. Our data from qPCR and immunofluorescence experiments corroborate information from scRNAseq, and bulk RNAseq analyses, which report a transcriptional landscape similar to what is exhibited at the protein level *in vivo*. Additionally, we show that the signature of this peritumoral microglia population is evident in 3 immunocompetent mouse models of HGG (GL261, KR158B, and RCAS from [8, 14]), confirming the translational accuracy of these markers and phenotypes.

### A Novel Transcriptional Signature Marks PGAM

While the above analyses define several specific molecular markers of PGAM, a more robust and comprehensive signature is needed to define PGAM across datasets and sampling modalities. 189 candidate transcripts for PGAM were identified between the DE analysis and literature review. To develop a faithful and spatially conscious transcriptional signature of PGAM, the selection was manually curated to render a list of 47 transcripts (Supplementary file 1) and their levels of expression were paneled across 3 anatomically annotated HGG datasets. Using myeloid cells from the Darmanis scRNAseq data in Figure 1, relative expression of the 47 proposed PGAM-specific genes, and the top 100 DE genes of tumor core GAM were plotted, and cells were grouped horizontally by tissue sample (core vs periphery) (Figure 3A). In line with the previous results, cells of the peripheral tissue samples express relatively high levels of PGAM genes. Signature scores were generated by calculating the average expression of PGAM and GAM gene lists for each cell. Plotting the signature scores indicated that cells of the peripheral tumor sample were enriched for PGAM gene signature, where cells of the tumor core were enriched for the GAM signature (Figure 3A).

**Figure 3.**
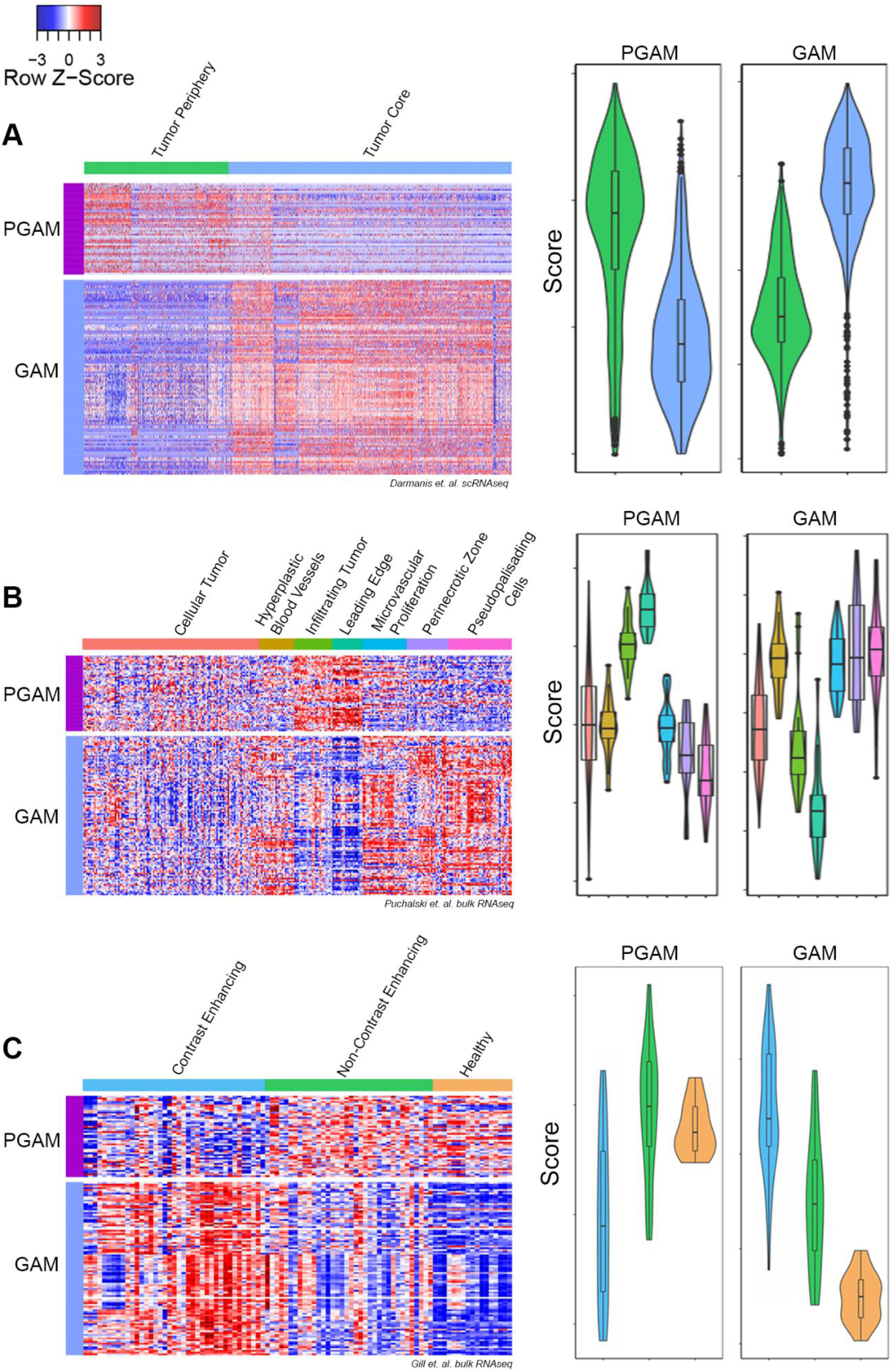
A novel PGAM gene signature. **A-C.** Left, heatmap of relative expression levels of PGAM and tumor core GAM genes, grouped horizontally by anatomical sample location. Rows are scaled by z score. Right, violin plots of signature scores generated by averaging PGAM or GAM gene expression across samples. Dataset derived from A. Darmais et al.; B. Puchalski et. al. (IVY GAP); C. Gill et. al. Y axes represent scores specific to each dataset, and are not on the same scale between gene signatures, or across datasets.

The same analysis was carried out on two additional datasets, both containing spatially annotated samples derived from human HGG. The Ivy Glioblastoma Atlas Project^39^ contains matched histologically annotated and bulk RNAseq samples derived from microdissection of fixed human specimens. The heatmap and gene scoring process revealed that the PGAM gene signature was exclusively enriched in leading edge and infiltrating tumor samples (Figure 3B). Conversely, tumor core GAM genes were expressed at relatively low levels in these same border regions. Importantly, both signatures were detected at moderate levels in cellular/core tumor samples. Using a third spatially annotated bulk RNAseq dataset, derived from MRI-guided biopsies of GB^40^, the PGAM gene signature was enriched in non-contrast enhancing/peritumoral samples, whereas contrast enhancing samples, representing the tumor core, correlated with GAM signature scores (Figure 3C). This result supports the spatial accuracy of the PGAM gene list, and suggests transcriptional/radiomic correlations.

The matched healthy reference tissue from patients in this dataset also shared positivity for PGAM-specific gene expression, which is in-line with the spatial overlap between CNS adjacent and peritumoral microglia. However, the presence of the PGAM signature score in cellular/core tumor samples and the wide range of scores in non-contrast enhancing/peritumoral samples suggest that this gene list captures microglia across the tumor/stroma interface. Further, this corroborates the observed activation spectrum reported in GAM-Mg^12,22,23^.

Thus, the PGAM-specific signature gene list, influenced by the expression of chemotactic/pro-inflammatory molecules, microglia-specific gene expression patterns, and border-associated transcripts reported in literature, faithfully identify PGAM in scRNAseq samples, and is recapitulated in bulk RNAseq with spatial correspondence. Our refined PGAM gene list proposed here expands on the existing 14 GAM-Mg specific list proposed by Muller et al^12^, and includes an additional 33 transcripts which comprehensively identify this novel subpopulation of GAM at the tumor/stroma interface (Supplemental File 1).

### Comparison of PGAM Across Glioma

There is a growing body of transcriptomic data, graciously made publicly available by research groups worldwide. We took advantage of this large pool of data to test the robustness of the PGAM signature across healthy and pathological samples. To capture the heterogeneity surrounding HGG, we collected data from primary and recurrent Grade IV GB, Grade III HGG, as well as IDH mutant astrocytoma and oligodenrocytoma^13,22,25,41,42^. It has been proposed that the observed activated GAM-Mg profiles reflect aging-related phenotypes of microglia^23^. Although these conclusions were made using reference samples presumed healthy, the samples were taken from patients afflicted with epilepsy, or undefined glioma/carcinoma^23^. After our quality control workflow was applied to these reference data, over 90% of microglia cells were removed (as mitochondrial genes made up the vast majority of detected transcripts, indicating compromised viability and data quality). To get a clearer transcriptional profile of healthy microglia, devoid of any pathology, and at single cell resolution, we sourced homeostatic microglia transcriptomes from the murine dataset provided by Huang et al^43^. Healthy murine-derived samples are not limited by tissue availability, and the PGAM signature is consistent in both human and mouse samples. In this dataset, scRNAseq was performed on FAC-sorted murine microglia under homeostatic conditions, as well as on repopulating microglia following a chemo-depletion regiment using the CSF1R inhibitor PLX3397^43^. This not only reflects resting CNS microglia, but also captures putative transcriptional programs of microglia during proliferation and local CNS repopulation conditions, thus spanning the physiology of healthy microglia.

We integrated these data to remove batch-related artifacts using the recently published tool LIGER^44^. LIGER applies an integrative non-negative matrix factorization (iNMF) approach to reduce batch and sample related artifacts, and captures relevant biological signal across data sets. The algorithm then builds on the strengths of NMF for clustering, dimensional reduction, and downstream analysis. LIGER asserts strengths in analyzing CNS tissue, as well as integrating data of different types, and across species^44^. It is also computationally efficient and scalable, compared to other scRNAseq integration methods^45^. LIGER was supplied with 48 samples and over 11,000 myeloid/microglia cells for integration and dimensional reduction. PGAM-labeled cells (annotated from the Darmanis dataset) separated from homeostatic microglia, repopulating microglia, as well as the bulk of tumor core GAM (Figure 4A). We identified key transcripts defining clusters using the feature loadings produced by the NMF reduction (Figure 4B). PGAM loaded on top PGAM signature genes, including *IPCEF1, CCL2, CCL3,* and *CCL4,* and also loaded on transcripts *DUSP6* and *A2M.* Two GAM sub-clusters were also identified. GAM subcluster 1 was enriched in a number of transcripts involved in active DNA remodeling and repair, including *ZBTB8A, ZNF850, POLH,* and *PDE4C,* whereas GAM subcluster 2 expressed various interferon-related genes, including *IFIT3, IFIT1, IFIT2,* and *ISG15.* Interestingly, GAM subcluster 1 was composed solely of cells from IDH mutant samples, while nearly all of GAM subcluster 2 was composed of cells from IDH wild-type patients. Along with the IDH status, the distinct transcripts may be indicative of unique metabolic phenotypes in these cells.

**Figure 4.**
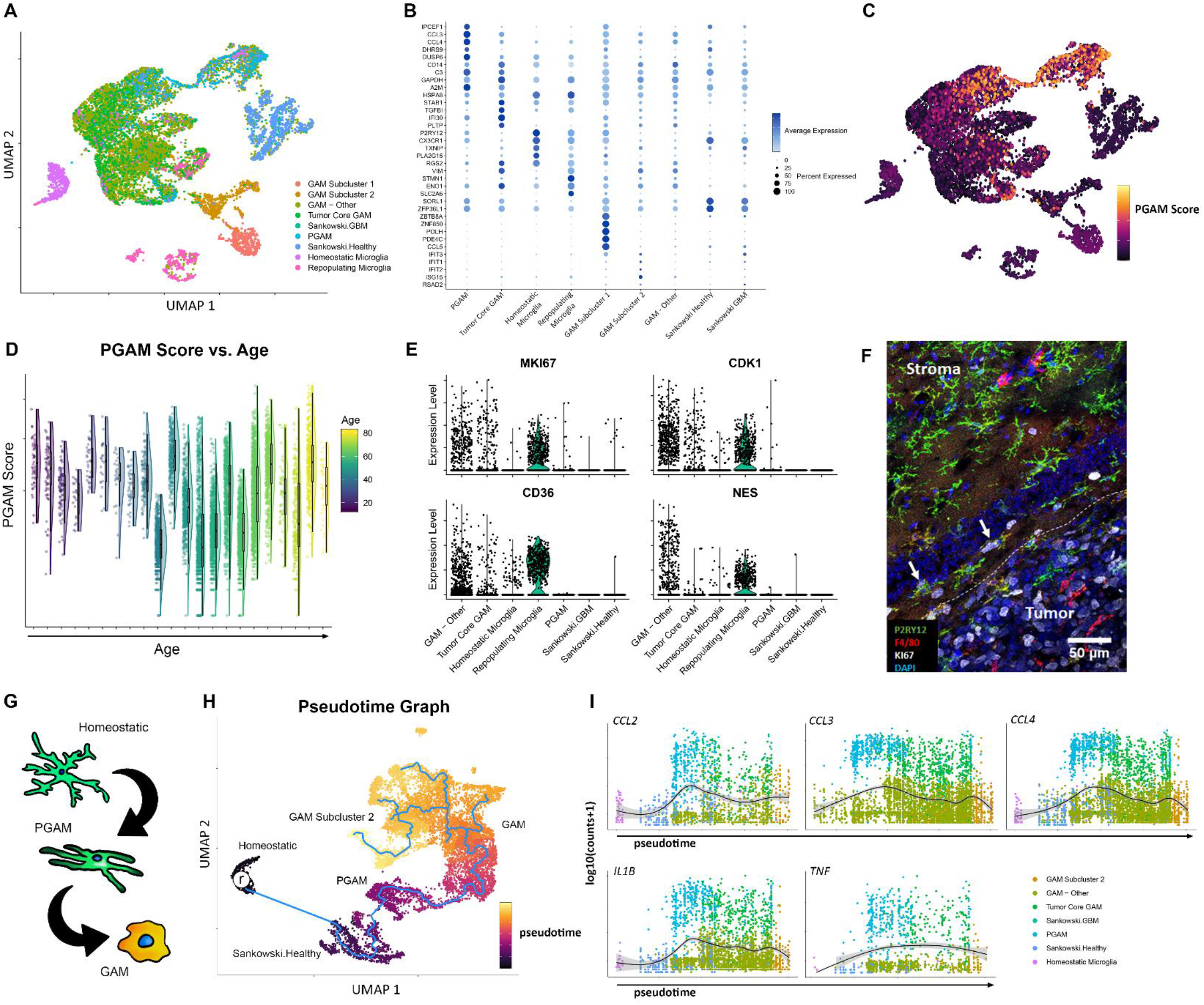
GAM across Health and Glioma. **A.** Myeloid cells from 47 samples were integrated using LIGER. UMAP was performed to plot the cells, which are colored by annotated populations. **B.** Feature loadings derived from LIGER were used to identify top transcripts contributing to cluster formation. The top 5 transcripts for each cluster are plotted, colored by the average expression across clusters, and sized by the percentage of cells expressing the transcripts. **C.** Cells form the integrated data were scored for PGAM gene signature, and the signature scores were plotted as an independent feature. **D.** PGAM scores for each cell were plotted against patient age. Raincloud plots shows the distribution of cell scores across each age bin. **E.** Violin plots of relative expression of the putative markers of repopulating microglia, across the annotated populations. GAM-Other contains all unidentified GAM and subclusters. **F.** Murine glioma sections were triple stained for Ki67, P2RY12, and F4/80. Ki67 positivity is primarily within the tumor core GAM, and neoplastic cells. White arrows mark triple positive cells. White dotted line marks tumor border. **G.** Outline of the predicted pattern of transcriptional evolution in peritumoral microglia. **H.** Pseudotime analysis of myeloid cells, following integration. ‘r’ marks selected homeostatic microglia acting as a root starting point for graph learning (blue line). Cells are colored by predicted pseudotime. **I.** Expression of chemotactic/proinflammatory genes are plotted in pseudotime. The expression is fitted with a smooth generalized linear model.

These data indicate that the PGAM transcriptional signature distinguishes GAM-Mg from microglia of the healthy brains at the single cell level, and is preserved across heterogenous disease states. When the scoring approach was used as in Figure 3, the PGAM signature faithfully identified these cells, and predicted nearby PGAM-like cells (Figure 4C). Notably, microglia of non-pathological sources were negative for the PGAM gene signature. We tested the hypothesis that the chemotactic/pro-inflammatory phenotype observed were related to aged microglia, and found no association between PGAM gene expression and patient age (Figure 4D). This finding suggests that the pro-inflammatory phenotype seen in GAM-Mg results from phenotypic changes at the leading edge niche.

As indicated by the dimensional reduction, repopulating microglia and PGAM form distinct clusters across the projected dimensions. Additionally, PGAM were negative for transcripts putatively marking the repopulating microglia population *MKI67, CDK1, CD36,* and *NES*^43^ (Figure 4E). To validate this finding, tissue sections from the murine HGG models were probed for the proliferation marker Ki67. Only a few double positive P2RY12^+^Ki67^+^ cells were present at the tumor/stroma interface, where most of the Ki67 signal was associated with tumor core GAM and neoplastic cells (Figure 4F). This demonstrates the ability of LIGER to accurately capture inherent differences between microglia populations across species. Collectively, these data indicate that the approach used here can accurately compare microglia between the healthy CNS parenchyma and glioma, and that PGAM transcriptional signatures coincide with a novel subpopulation of chemotactic and proinflammatory GAM that is maintained across heterogeneous disease. This integrated dataset serves as a resource collectively describing microglia across health and glioma. The data will be made available, upon publication.

### PGAM constitute an intermediate between healthy microglia and GAM

To further elucidate the cellular origins and dynamics of PGAM, additional bioinformatic analysis was employed using the pseudotime learning algorithm provided by Monocle3^46,47^. To account for phenotypic variability, we removed cells of the GAM subcluster 1 and repopulating microglia groups. This allowed a more relevant organization of clusters. Remaining cells were re-clustered with UMAP. Cells of the homeostatic microglia cluster were selected as a physiological “root”, representing a starting point from which all microglia/myeloid cells could transcriptionally deviate. The pseudotime trajectory moves from homeostatic microglia, to the human derived healthy reference microglia, and then to the PGAM cluster (Figures 4G, 4H). This organization indicates that both sources of reference microglia serve as a precursor to PGAM transcriptional states. The trajectory continues into the bulk GAM cluster, where bifurcations occur between GAM subcluster 2, and other potentially distinct GAM sub-states. In concordance with the chemotactic/proinflammatory transcripts identified in Figure 2, we mapped the expression of these genes in pseudotime, and found similar expressional patterns, with the PGAM population having peak expression levels of *CCL2, CCL3, CCL4, IL1B,* and *TNF* (Figure 4I).

The results from pseudotime analysis give insights on the observed activation spectrum of GAM-Mg, and potentially implicate PGAM as an intermediate microglia population at the tumor/stroma interface. The current analysis, however, does not delineate directionality of PGAM transition; that is, whether or not these cells are drawn into the tumor, or are ‘casualties’ of the expanding margin. Future work incorporating a temporal dimension of PGAM behavior would more equivocally assign such behaviors.

### PGAM Enrichment Correlates with MRI, Immune Signatures, and Poor Prognosis in GB

We next investigated the relationship between the PGAM gene signature, and clinical covariates of disease. REMBRANDT is as a collection of clinical, transcriptional, and radiomic data, which serve to enhance investigation and interoperability of brain neoplasias^48^. 28 patients who had both radiomic and transcriptomic data were scored for PGAM enrichment, and split into either high or low groups based on median gene signature score. Using the 30 reported radiomic features, a Chi-squared test was performed between the two PGAM groups, which revealed several statistically significant associations between PGAM enrichment and poor enhancement quality of HGG lesions (Supplementary File 1). Notably, high levels of PGAM significantly correlated with absence of contrast-enhancing tumor regions and margin in patients, and a strong, but not significant, increase in non-contrast enhancing proportions of tumor (Figure 5A, Supplemental file 1). This finding corroborates the earlier results, in which PGAM scoring was greater in MRI-guided biopsies of non-contrast enhancing samples, which represented the peritumoral space^40^ (Figure 3C). Together these joint bulk RNAseq and radiomic analyses indicate that peritumoral microglia, described by a unique gene signature, can be associated with MRI contrast enhancing features in HGG.

**Figure 5.**
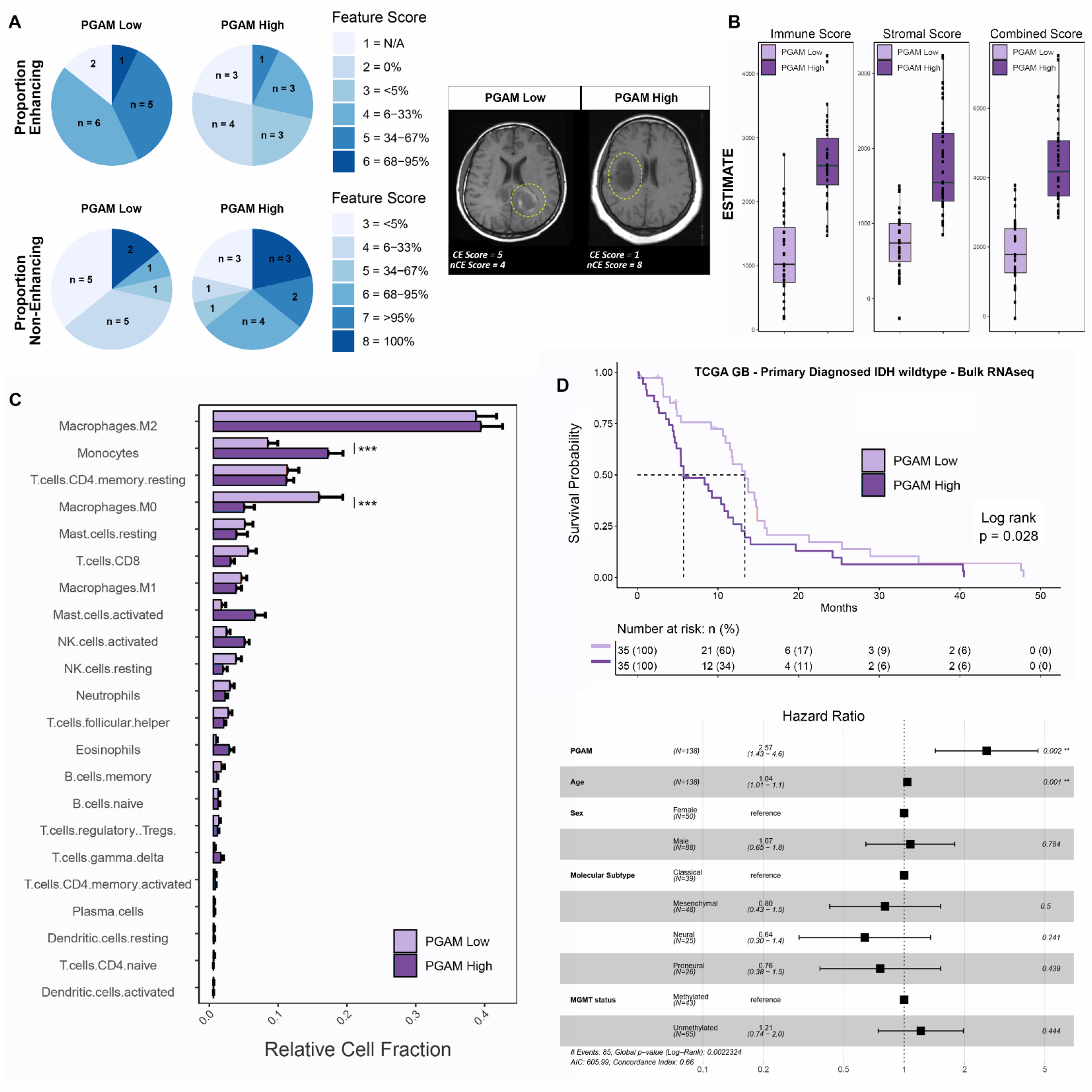
PGAM correlate with clinical covariates of GB. **A.** Patients were split into high/low PGAM signature scoring groups. Left, pie charts of feature scores, by PGAM group. n represents the number of patients with each feature score, and are colored by increasing feature score severity. Proportion of enhancing, and non-contrast enhancing tumor regions are shown. Right, representative MRI images from each PGAM group, and their corresponding feature score. CE = contrast enhancing, nCE = non-contrast enhancing. Yellow dotted line demarks primary lesion. n = 14 per group. **B-D.** Primary diagnosed GB IDH wildtype patients from TCGA-GB cohort were scored for PGAM gene signature, and split into high/low scoring groups. n = 35 per group. **B.** Boxplot of ESTIMATE immune and stromal scores, stratified by PGAM signature scoring.**C.** Patient transcriptomes from PGAM high/low groups were analyzed by CIBERSORT. Plotted are the predicted relative fractions of 22 immune cell subpopulations ±SEM. Two-way ANOVA with Tukey’s post hoc comparison. p<0.001***. **D.** Top, Kaplan-Meier survival curve, stratified by PGAM group. Dotted line marks median survival. MS PGAM low = 5.7, MS PGAM high = 13.3. Log rank p val = 0.028. Bottom, Cox proportional hazards regression model was conducted, using covariates of survival. Forrest plot of the resulting corrected hazard ratios, following regression.

As radiomic results showed clear association between PGAM enrichment and glioma, we examined a possible connection between these pro-inflammatory peritumoral microglia, and immune signatures of the TME. Using the widely regarded TCGA-GB cohort^49^, 138 primary diagnosed and IDH wildtype patients were scored for PGAM enrichment. Patients with highest PGAM gene expression exhibited markedly higher ESTIMATE scores for immune and stromal signatures^50^ (Figure 5B), likely reflecting high numbers of peritumoral microglia and other glial cell involvement at the tumor leading edge, and an association with systemic immune cell infiltrates. Using the CIBERSORT *in silico* deconvolution algorithm, immune cell populations present in these samples were characterized^51^. CIBERSORT uses genetic profiles of 22 immune cell populations to derive relative and absolute cellular fractions of tumor biopsy transcriptomic data. Of the 22 paneled cell types, infiltrating monocytes were enriched in highly scoring PGAM patients, while resting macrophage populations were enriched in PGAM-low samples (Figure 5C). As PGAM are defined by elevated levels of *CCL2, CCL3,* and *CCL4* expression, these results highlight this important signaling axis in GB^30-32^, where PGAM may represent a potent source of monocyte/macrophage recruitment to the TME.

GAM recruitment negatively impacts patient survival in HGG, particularly by affording the growing neoplasm with tumor promoting properties, and sculpting an immunosuppressive TME^5,7^. The currently prevalent hypothesis is that polarization of GAM to a pro-inflammatory phenotype may facilitate tumor abrogation^18,52-54^. To examine if the chemotactic and pro-inflammatory phenotype of PGAM that correlates with monocyte recruitment also correlates with GB outcomes, Kaplan-Meier survival analysis was performed with primary diagnosed, IDH-wildtype GB patients from TCGA, and stratified by PGAM gene signature scoring (Figure 5D). A survival disadvantage was observed in patients scoring highly for PGAM genes (median survival ~5.7 months, vs ~13.3 median survival of PGAM low group). Multivariate cox-proportional hazards analysis was performed to regress out potentially confounding factors of survival in this patient cohort. Against several diagnostic and investigational clinical covariates of GB, PGAM signature scoring was significantly associated with poorer overall survival (Figure 5D, bottom).

## 3. Discussion

Over the past decade there has been great focus on modulating the TME of solid tumors to deplete neoplastic lesions of their support and foster therapeutic sensitivity. Success in the immunotherapeutic space has supported this effort, however, there have been limitations to this strategy, as the TME is not fully characterized. Specifically for GB, although many immunotherapy-based clinical trials are underway, little success in extending survival has been achieved. The progression and recurrence of GB have been attributed to the highly invasive phenotype of neoplastic cells, their chemo- and radio-therapy resistance, and the support of the CNS TME. As such, myeloid cells of the TME have become a focus of investigation. Description of local TME niches will help decipher the roles of these cells in disease progression and may uncover novel therapeutic targets.

Using several orthogonal approaches, we comprehensively and methodically characterize a previously undescribed microglia population in GB, which comprise a subpopulation of GAM. We define PGAM as peritumoral microglia, marked by chemotactic and pro-inflammatory gene enrichment, dynamic P2YR12 expression, and spatially correlate with the leading edge of glioma. The spatial context provided with changes in P2RY12 expression may reflect the transition of nearby resting/homeostatic microglia adopting traditional GAM phenotypes, as they become incorporated into the neoplastic lesion.

With our results, we produce a novel gene signature that serves to define PGAM in both scRNAseq and bulk RNAseq data, and show that it can unmask these cells in both human and mouse derived samples. This list could be applied in future studies to identify fractions of PGAM in HGG tissue samples, an understudied myeloid group with consequences yet to be resolved. The PGAM signature associated with clinical and radiomic features of HGG. As MRI is the major method of patient follow-up and used to detect HGG progression, integrating this transcriptional signature with radiomics may also prove beneficial in the clinical setting.

Potent inflammatory and chemotactic molecules make up the PGAM gene signature. A shift towards pro-inflammatory GAM polarization is traditionally thought to be beneficial in cancer^53-55^. However, we caution that high levels of pro-inflammatory chemokines and cytokines may need to be carefully examined. If a primary function of the innate immune system is to further recruit innate and adaptive immune cells, we predict that unchecked recruitment of inflammatory monocytes to the neoplastic site seals the fate of the TME, where the cells ultimately become subverted into adopting pro-tumorigenic and immunosuppressive programs^2,7^. In line with this idea, a recent report has demonstrated that TLR2 activation of myeloid cells at the glioma border confers a pro-inflammatory and pro-tumorigenic shift, which ultimately prevents antigen presentation and supports tumor growth^56^. Thus, in agreement with other studies, we support adopting the idea of ‘myeloid checkpoint inhibition’^54,57,58^, which describes strategies aiming to modulate pro-tumorigenic functions of GAM. Such strategies would halt the effector and pro-tumorigenic functions of macrophages, and thus attenuate the immunosuppressive and tumor-supportive functions of the TME. In the presented analysis several targets have been identified for further investigation, and align with this idea.

There is accumulating evidence to suggest that P2RY12 dynamically regulates glioma/microglial interactions and modulates the proliferative capacity of glioma cells. A series of elegant studies by Chia et al. demonstrated that glioma cells adapt neuron-microglial signaling mechanisms to fuel their own proliferation^59^. Akt1^+^ preneoplastic cells exhibit higher cytosolic calcium levels via activation of ionotropic glutamate NMDA receptors. Increased cytosolic calcium levels trigger ATP release, which activates P2RY12 receptors on microglia, and leads to microglial process extension and an increase in Akt1^+^/microglial cell interactions^59^. NMDA antagonism and CRISPR/Cas9-mediated deletion of P2RY12 reduced interactions, and significantly perturbed Akt1^+^ cell proliferation. Antagonizing P2RY12 receptors on microglia may therefore abolish these pro-proliferative interactions. P2RY12-ATP driven process extensions in microglia have also been implicated in epilepsy. During periods of hyperactivity of neuronal circuits during epileptic events, glutamate release activates post-synaptic NMDA and AMPA receptors with downstream events leading to increased ATP efflux from the post-synaptic cell; the ATP efflux activates microglial process extension via P2RY12 toward the neurons releasing ATP^60^. Again, when P2RY12 was ablated, microglia were unable to respond to the hyperactivity, and mice exhibited severe epileptic seizures^60^. Microglia P2RY12 signaling is also linked to aberrant seizure-induced neurogenesis^61^. As recent studies have outlined substantial neuronal-neoplastic communication in GB^62,63^, the presence of non-uniform populations of P2RY12^+^F4/80^High^ peritumoral microglia, and mature neurons at the glioma leading edge suggests that a potential tripartite system exists including neurons, glioma, and myeloid cells. Future work should further detail P2RY12 molecular involvement in HGG.

Microglial transcriptomes are known to be influenced by aging, and P2RY12 has been shown to be significantly downregulated with age, however, direct phenotypic effects of these changes have not been adequately studied^64-66^. As age is a primary clinical covariate in HGG, the patterns and consequences of microglial P2RY12 expression and resulting GAM phenotypes need further investigation^67,68^. We did not observe any correlation between the expression of the PGAM gene signature (which includes P2RY12) and aging, supporting the idea that rapid changes in expression results from underlying shifts in response to the TME, rather than as a function of age.

One of the main pro-inflammatory cytokines associated with the PGAM signature is CCL2. The consequences of CCL2-mediated monocyte recruitment in solid tumors, such as glioma, has recently surfaced, and an increased number of infiltrating monocytes has repeatedly been shown to be detrimental to HGG outcome. Our results potentially place PGAM at the crux of this signaling axis. Glioma cells secrete CCL20 which induces neighboring myeloid cells to secrete CCL2, thus aiding to attract peripheral monocytes and T regulatory cells through CCR2 and CCR4, respectively^32^. Peripheral monocytes can differentiate into myeloid-derived suppressor cells (MDSCs) and, along with immunosuppressive T_regs_, inhibit cytotoxic T-cell responses^32^. The observation that CCL2 mediates recruitment of multiple immunosuppressive cell types is supported by the finding that dual CCR2 antagonism and PDL1 checkpoint blockade synergistically prolong the survival of glioma-bearing mice^30^. While such results are encouraging, antagonizing CCR2 to slow the accumulation of immunosuppressive MDSCs does little to prevent the accumulation of CCR4^+^ T_regs_, which suggests that sequestering CCL2 may be a superior treatment modality.

In a clinical study of diabetic nephropathy marked by infiltration of pro-inflammatory monocytes, CCL2 antagonism was accomplished by treating patients with the RNA aptamer emapticap pegol (NOX-E36)^69^. Administration of the CCL2-sequestering ligand was well-tolerated and reduced albuminuria, a hallmark of diabetic nephropathy. Sequestering CCL2 in the bloodstream could also circumvent the common obstacle of designing drugs that can penetrate the blood-brain barrier.

## Conclusions

In sum, a subpopulation of microglia of the peritumoral space can be uniquely identified, and may influence disease outcomes by recruiting systemic monocytes to the TME. The possibility to detect these outcomes using a combination of radiomic and transcriptomic approaches are an attractive option, where new applications integrating genomic and clinical data are being pursued. Profiling myeloid cells in further detail may uncover targets and signaling pathways, which could be exploited in solid tumor malignancies, such as GB.

Our analysis identifies a subset of microglia present in glioma invading margins, and suggests a role for them in tumor promotion. Potential targets could be leveraged to modify pro-tumorigenic functionality of glioma-associated microglia.

## 5. Methods

### Immunocompetent glioma cells lines

GL261 cells were obtained from Dr. Michael Lim’s lab, and are derived from a chemically induced C57BL6 murine astroglioma^27^. KR158 cells were obtained from Drs. Tyler Jacks’ and Behnam Badie’s labs, and are genetically engineered *Nf1/Tp53* mutants^28^. Cells were cultured in Dulbecco’s Modified Eagle’s Medium, 10% serum, 1% antibiotic, and 1% sodium pyruvate, at 37°C 5% CO2.

### Murine glioma model

Orthotopic gliomas were established in adult C57Bl6 mice. Briefly, mice were anesthetized, a midline incision was made on the scalp, and a small burr hole was drilled in the skull at stereotactic coordinates of bregma, −1 mm anteroposterior and +2 mm mediolateral. 1μL of 3×10^4^ cells were injected over 2 minutes at a depth of 3 mm. After completion of the procedure, the incision was sutured and the mice were placed on a heated surface until fully recovered from anesthesia.

### Immunofluorescent staining

Whole brains were collected from animals on days 15 and 21 post tumor implantation. Animals were transcardially perfused with 4% paraformaldehyde, brains were dissected, and post fixed overnight, cryopreserved in 30% sucrose for 24 hours, and then embedded for cryosectioning. 20μm tissue sections were collected on Superfrost™ plus microscope slides, and stored at −80°C until use. For immunostaining, sections were brought to room temperature, and washed in PBS 3x. Slides were then blocked with 5% serum, 0.3% Triton X-100 in PBS for 1 hour at room temperature. Primary antibody was incubated overnight at 4°C. Slides were washed with PBS-T, and secondary antibody was incubated 1 hour at room temperature. Sliders were again washed with PBS-T, and mounted with Fluoromount-G™ (Thermo, #00-4958-02). See supplemental methods table for dilutions. Confocal imaging was performed using the Lecia Sp8-x system, with white light and argon lasers.

### RNA isolation, cDNA synthesis, and quantitative real-time PCR

Whole brains were collected from animals on day 21 post-tumor implantation. Microdissection was performed to sample ~100mg of tissue from the tumor core, peritumoral tissue, and contralateral tissue. Total RNA was collected with TRIzol™ (Thermo, #15596026) following the manufacturer’s protocol. cDNA libraries were created using the High Capacity cDNA Reverse Transcriptase Kit (Applied Biosystems, #4368814). Real-time quantitative PCR was performed using a StepOnePlus™ thermocycler. A cutoff of 35 Ct was applied, and ΔCt values were calculated by normalizing to housekeeping gene GAPDH. Given that some transcripts were not detected in healthy tissue, ΔΔCt was not calculated. ΔCt values were log transformed and used for analysis.

### Bioinformatic Analysis of publicly available datasets

#### Bulk RNAseq

For all bulk RNAseq samples, data were downloaded from the GEO, and normalized using log2(FPKM+1). For murine analysis of pro-inflammatory chemokines, normalized datasets were merged by common gene ids, and processed for batch removal using Combat^70^. For other data and analysis, heatmaps were created using a modified function of heatmap3.

#### scRNAseq

Raw counts matrices were downloaded from GEO, and expression objects were created using Seurat^71,72^. Quality control was performed, removing cells with abnormally high or low feature counts, percent mitochondrial gene expression < =10%, and percent ribosomal gene expression < =25%. In accordance with [^23^], cells were also removed based on percent *KCNQ1OT1* expression < =2%. Following merging of samples, features expressed in a minimum of 40 cells were retained. After QC was applied, samples containing less than 20 cells were omitted. Myeloid cells were selected from datasets based on *PTPRC, FCGR3A, CD14,* as well as other canonical monocyte marker expression. To integrate the murine dataset, mouse transcripts were converted to human transcripts by selecting for orthologs, derived from biomaRt^73,74^ gene symbol lists. Only direct orthologs, or mouse transcripts mapping to unique human transcripts were retained.

Integration of single cell data was performed across patient samples, using the following parameters for LIGER, and SeuratWrappers provided functions: variable features = 3000, k = 13, lambda = 8. UMAP was performed with the following hyperparameters: n.neighbors = 40, min.dist = 0.6. iNMF projections were passed to Monocle3, and pseudotime graph was learned using the default parameters.

#### Survival Analysis

transcriptomic and clinical data were downloaded from TCGA GB cohort using TCGABiolinks. Healthy, recurrent, and IDH mutant samples were omitted. Patients were scored for gene signatures, and Kaplan-Meier survival analysis was performed. Multivariate cox regression analysis was conducted, using age, sex, MGMT status, and molecular subtype as covariates.

## Declarations

### Ethics approval

All work described in this manuscript was approved by the Stony Brook University IACUC committee (Protocol 246938).

### Consent for publication

All authors are in agreement with the content of this manuscript.

### Availability of data and materials

Data will be made available, upon acceptance for publication.

### Authors’ contributions

MDC designed, performed and analyzed experiments, wrote and edited drafts of the manuscript; KO analyzed experiments and edited drafts of the manuscript; MM, DR, NS performed and analyzed experiments, and edited drafts of the manuscript; RAM analyzed experiments and edited drafts of the manuscript; SET designed and analyzed experiments, wrote and edited drafts of the manuscript.

## Acknowledgements

We would like to thank members of the Tsirka and Moffitt labs for helpful suggestions along the way. Artwork credit Lucie Chrastecka.

---

**Supplemental Figure 1.**
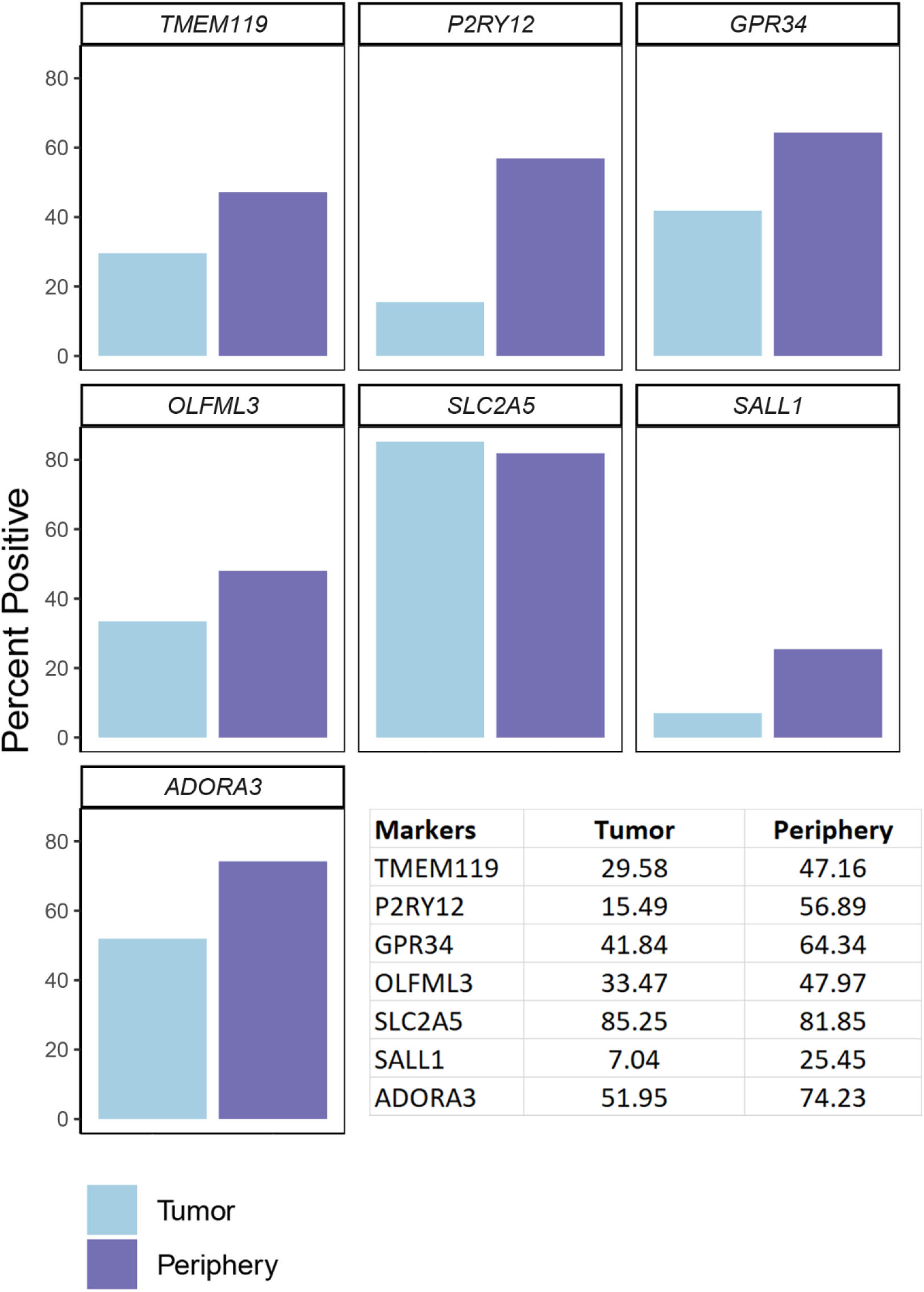
Percent of cells expressing microglia core transcripts, plotted across tumor compartment in the Darmanis et al dataset.

**Supplemental Figure 2.**
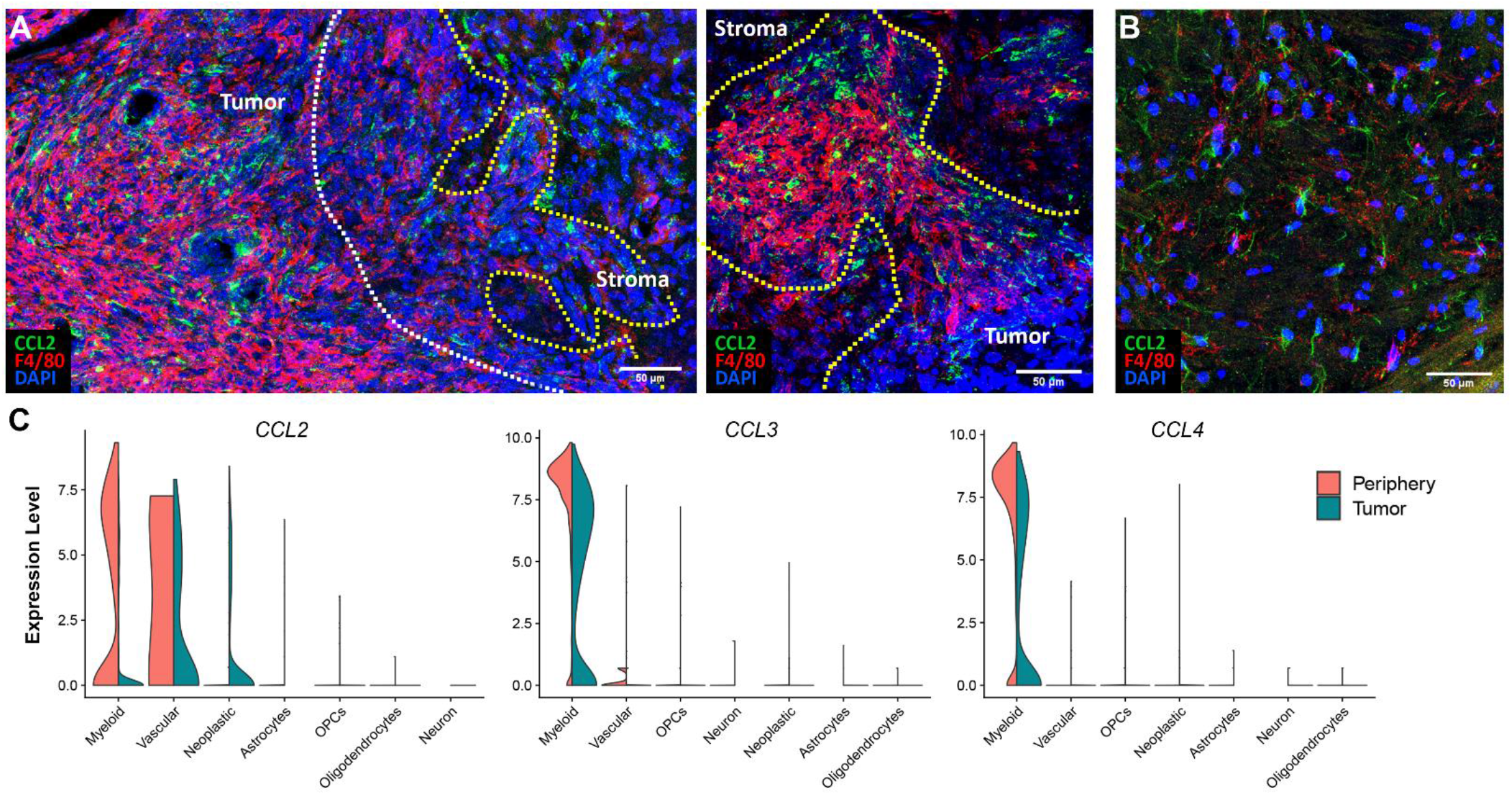
**A.** CCL2^+^ cells populate peritumoral space, and infiltrating zones, in KR158 glioma samples. White dotted line indicates bulk tumor border. Yellow dotted line indicates infiltrating zones. n =3. **B.** Representative image of CCL2^+^F4/80^Low^ cells in the contralateral hemisphere of male mouse harboring glioma. **C.** Violin plots showing relative expression of *CCL2, CCL3,* and *CCL4* in all CNS cell populations, from Darmanis et. al single cell dataset. Plots are split and colored by sample location.

**Supplemental Figure 3.**
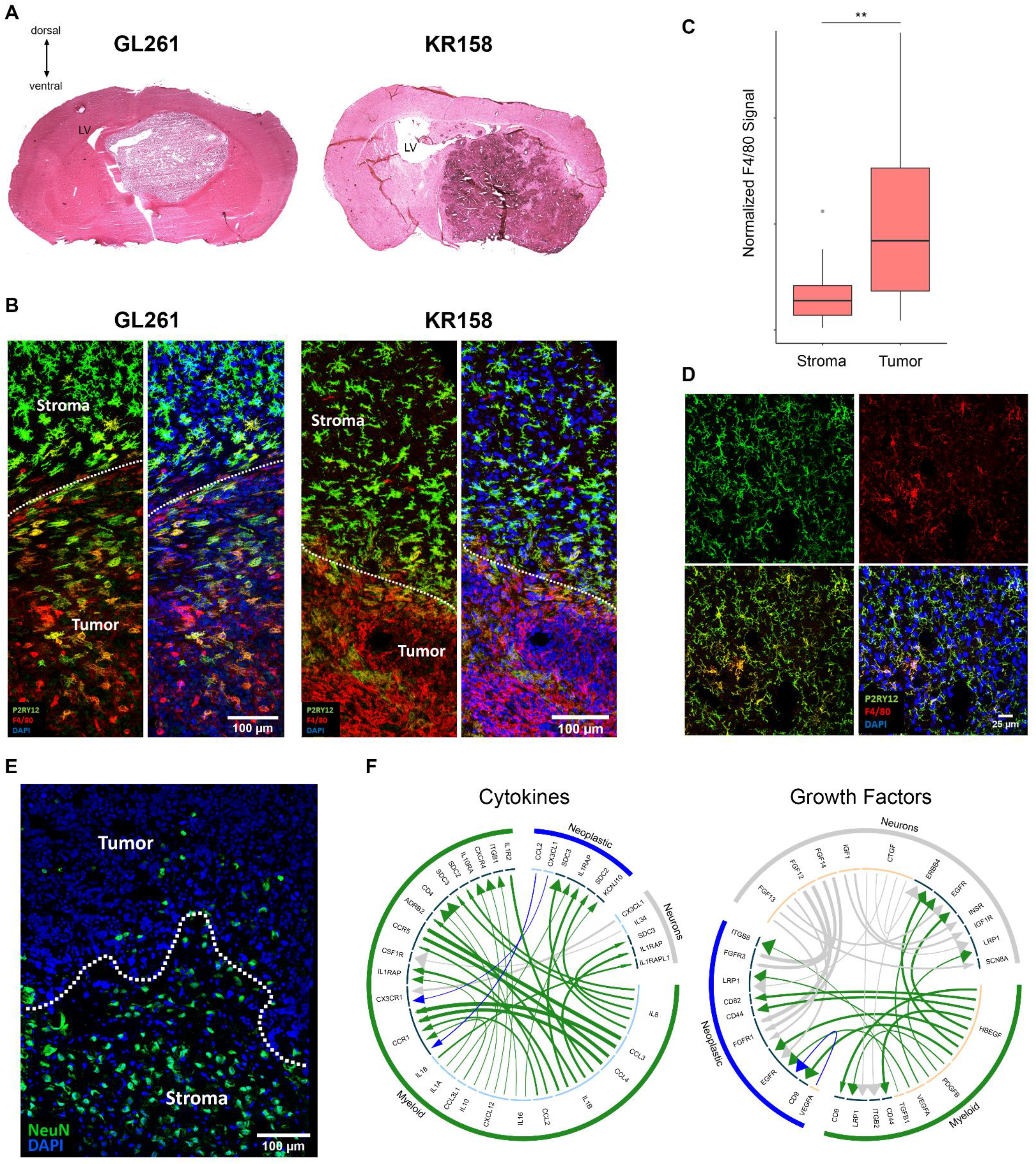
**A.** Representative H&E images of GL261 and KR158 murine gliomas, at peak disease. **B.** P2RY12 F4/80 co-staining of GL261 gliomas on day 15, representing progressive disease (left), and KR158B tumors at peak disease (right). n = 2-3. **C.** Spatial mapping of F4/80 expression, from figure 2G. **D.** Healthy microglia from contralateral hemispheres show P2RY12^+^F4/80^Low^ expressional patterns. **E.** NeuN staining marks mature neuron cell bodies at the tumor/stroma interface in GL261 gliomas. **F.** iTALK receptor-ligand pairs are predicted based on relative expression, between myeloid and neoplastic cells. Arrows are colored by cellular population. Arrow lines are scaled by ligand expression, arrow heads are scaled by receptor expression. Inner circle: light blue denotes ligands; dark blue denotes receptor.

